# Variation in self-compatibility among genotypes and across ontogeny in a self-fertilizing vertebrate

**DOI:** 10.1101/2021.02.22.432320

**Authors:** Jennifer D. Gresham, Anna Clark, Chloe M.T. Keck, Alexis E. Longmire, Abye E. Nelson, Haylee M. Quertermous, Ashley B. White, Ryan L. Earley

## Abstract

Mixed mating strategies offer the benefits of both self-fertilizing one’s own eggs (selfing) and outcrossing, while limiting the costs of both methods. The economics of mixed mating is further determined by individual self-compatibility. In gynodioecious (hermaphrodites, females) and androdioecious (hermaphrodites, males) species, the level of self-compatibility of the hermaphrodites also acts as a selection pressure on the fitness of the other sex. Mangrove rivulus fish populations are comprised of selfing hermaphrodites and males that result from hermaphrodites changing sex. Although hermaphrodites overwhelmingly reproduce through internal selfing, they occasionally oviposit unfertilized eggs. Males can externally fertilize these eggs. Here, we reveal that fecundity and self-compatibility varies within individuals across ontogeny until about 365 days post hatch, and among individuals derived from lineages that vary in their propensity to change sex. Hermaphrodites from high sex changing lineages were significantly less fecund and self-compatible than hermaphrodites from low sex changing lineages. These differences in self-compatibility and fecundity have the potential to drive evolutionary changes on mating strategy and the fitness of males in populations of the mangrove rivulus. This study also illustrates the importance of including lineage variation when estimating the costs and benefits of mixed mating strategies.

## Introduction

As a reproductive strategy, self-fertilization has many benefits. It overcomes the “cost of males,” wherein only one half of the population can produce offspring compared to the entire population in asexual or self-fertilizing species (Maynard Smith 1971, 1978). Self-fertilization also assures reproduction when pollinators and/or mates are scarce (Darwin 1876; Baker 1955; Stebbins 1957; Pannell et al. 2015), eliminates physical damage associated with mating and/or sexually transmitted parasites or pathogens (Fowler and Partridge 1989; Clutton-Brock and Parker 1995), and maintains well-adapted genotypes in stable environments (Allard et al. 1972; Dolgin et al. 2007). However, selection for selfing is often counteracted by selection for outcrossing. Benefits of outcrossing include combatting inbreeding depression in two ways: i) preventing accumulation of deleterious alleles in single genotypes, and ii) maintaining heterozygotes that may be more fit than homozygous individuals (Muller 1932). Ultimately, outcrossing represents a bet hedging strategy because it increases the variability of offspring fitness against a changing abiotic environment, or against coevolving parasites, predators and competitors (Muller 1932; Bell 1988; Lively 2010; Morran et al. 2011; Hartfield and Keightley 2012).

Many plant and invertebrate species have evolved a mixed mating strategy of selfing and outcrossing. Mixed mating allows individuals and populations to reap the rewards of both strategies as biotic and abiotic factors change the fitness landscape across space and time. Because the fitness landscape is variable, it would be expected that self-compatibility also would vary both within and among populations due to changes in the magnitude and/or type of selection acting for and against selfing and outcrossing (Escobar et al. 2011). Plant populations that are often mate- and/or pollinator-limited at the range edge or in ephemeral habitats have increased self-compatibility compared to other populations, as reported for the small angiosperm *Leavenworthia alabamica* (Busch 2005). Populations found near the center of the range and in more permanent habitats are much more likely to be self-incompatible (Busch 2005). When among-individual variation in self-compatibility is heritable, it provides the genetic and phenotypic background on which selection can drive evolutionary change in mating strategies. There are multiple examples of heritable variation in the self-compatibility of plants, including in the creeping bellflower, *Campanula rapunculoide* (Stephenson et al. 2000), and the Carolina horsenettle *Solanum carolinense* (Travers et al. 2004). Genetic and phenotypic variation in self-compatibility, along with environmental variability, can thus drive variation in mating strategy, as they alter the economics of selfing and outcrossing.

In mixed mating populations, self-compatibility can also vary across ontogeny, perhaps as a plastic response to environmental conditions. Individuals in populations that have higher rates of outcrossing often become more self-compatible with increasing age. This is often the result of selection favoring a “waiting” strategy for outcrossing opportunities when individuals are young, presumably to derive the benefits of variable and more fit progeny but also to assure some reproduction even if mates and/or pollinators are limited. This has been documented in plants (*Campanula rapunculoide*; Stephenson, Good, and Vogler 2000, *Solanum carolinense;* Travers, Mena-Ali, and Stephenson 2004), and snails (*Physa acute*; Tsitrone, Jarne, and David 2003), but few other self-fertilizing species. Populations can also evolve different mating strategies based on age structure. If younger individuals are more populous, and mating opportunities are plentiful enough to make selfing unnecessary, age-dependent changes in self-compatibility may be lost so that little or no selfing occurs. Alternatively, if populations consist predominantly of older individuals, selfing may continue to be an evolutionary stable component of a mixed mating strategy.

In androdioecious species, in which populations consist of hermaphrodites and males, variation in self-compatibility should also result in variation in the proportion of males within a population. Populations consisting of individuals with lower self-compatibility should maintain a higher proportion of males, as the “cost of males” is decreased by more outcrossing opportunities. Hypotheses concerning the evolution and variation of self-compatibility and mixed mating have been examined in many plant and invertebrate systems (Busch 2005; Winn et al. 2011; Wright et al. 2013), but not in a vertebrate species. Here we present evidence for individual variation in self-compatibility in one of only two vertebrates known to utilize a mixed mating strategy, the mangrove rivulus fish (*Kryptolebias marmoratus*).

The mangrove rivulus (hereafter “rivulus”), is a small euryhaline killifish that inhabits high elevation mangrove forests of Florida, the Bahamas, and Central America. Populations consist primarily of self-fertilizing hermaphrodites with varying proportions of males (no females). Rates of outcrossing and selfing also vary among populations and correlate with the proportion of males in the population; outcrossing rates increase as the proportion of males increase (Mackiewicz et al. 2006b; Tatarenkov et al. 2009). While self-fertilization happens internally, males can only achieve reproductive success if/when hermaphrodites lay unfertilized eggs, and there is no evidence that hermaphrodites outcross with each other (Furness et al. 2015). Rivulus males result from sex change, after which selfing hermaphrodites that make the sexual transition become obligate outcrossers. There also appears to be genetic variation for the propensity to change sex (Gresham et al. 2020, Turner 2006). Hermaphrodites fertilize the vast majority of their own eggs (> 94%, Harrington 1971, personal observations), giving males very limited opportunities for reproductive success. Previously, it was reported that while individuals that change sex to male are significantly more likely to survive environmental challenges than those that remain hermaphrodite, they are much less likely to lay any eggs before sex change occurs (only 2 out of 180, Gresham et al. 2020). This presents the problem of how individuals that change sex might successfully pass their sex-changing alleles on to the next generation.

We hypothesized that the proportion of fertilized eggs (a measure of self-compatibility) would change with age and/or the propensity of a given lineage to change sex. First, we predicted that self-compatibility would increase with age. Cole and Noakes (1997) reported histological evidence that ovarian tissue matures before spermatogenic tissue. We predicted that there would be a period immediately after sexual maturity where oviposition of unfertilized eggs would be more likely. Second, we predicted that overall fecundity and/or self-compatibility would be greater in lineages that are more likely to change sex. This prediction is based on two competing ideas about how sex-changing alleles are successfully maintained in rivulus populations: i) selection should favor individuals in high sex changing lineages to be very fecund and self-compatible prior to changing sex, so as to compensate for any lost reproductive success after the transition, and/or ii) selection should favor individuals derived from lineages that change sex frequently to be highly fecund and self-compatible so that the siblings of sex changers recoup the lost reproductive success of their “brothers.”

## Methods

We designed and executed an experiment to measure the fecundity and self-compatibility of 213 individual fish representing 100 genetically distinct lineages (determined via microsatellite genotyping using 32 loci; (Mackiewicz et al. 2006a; Tatarenkov et al. 2010). We calculated the propensity to change sex for each of the 100 lineages as the proportion of individuals in each lineage that transition from hermaphrodite to male under common garden conditions (Gresham et al. 2020), which ranged from 0 - 68.6%. All experimental fish were raised from hatching in individual plastic containers (Rubbermaid^®^ Take-a-long Deep Squares) filled with approximately 200 mL 25‰ synthetic saltwater (prepared with Instant Ocean^®^ sea salt and aged tap water) until 30 days post hatching (dph), at which time water volume was increased to 600 mL (25‰). At 60 dph, we suspended a large marble-sized ball of egg-laying substrate (Poly-fil^®^) from one top corner of the container. Beginning at 67 dph, each container was checked weekly for eggs in the substrate, around the top edge of the container, and on the bottom of the container. Fish were checked weekly for eggs until they changed sex to male or were 365-371 dph, ensuring that each remaining hermaphrodite was at least 365 dph and had been checked weekly 43 times.

Eggs were collected into 59 mL plastic cups (Fabri-Kal^®^ Greenware GXL250PC) filled with approximately 6 mL 25‰ synthetic saltwater. Eggs were immediately counted and viewed under a stereomicroscope. Eggs were photographed and scored as either fertilized, unfertilized, or inviable. Fertilized and unfertilized eggs were identified by the presence or absence of a perivitelline space, respectively. Inviable eggs were identified by their marked discoloration and opaque appearance (Figure 1). Unfertilized and inviable eggs were removed from the cup and discarded. Fertilized eggs were left in the cups, and water was replaced every 7 days. Cups were checked every day for hatchlings. Hatchlings from the experimental animals were then scored as alive or dead, with live hatchlings being euthanized in 4°C water. Experimental fish were also observed for signs of sex change each week when the tubs were checked for eggs, as described in (Scarsella et al. 2018); orange freckles or orange skin is the most reliable external character of sex change. Individuals that changed sex (35 of the 213) were separated from the others and the date and age recorded. All experimental animals were isolated from the general colony for the duration of the experiment, kept on a 12h light: 12 h dark photoperiod, and fed a 4 mL suspension of brine shrimp (*Artemia*) nauplii (∼2000 shrimp) six days per week. Room temperature was also maintained at 26.45 ± 1.6 °C (mean ± SEM). The University of Alabama Institutional Animal Care and Use Committee approved all procedures (IACUC Protocol #18-10-1644).

**Figure 1.**
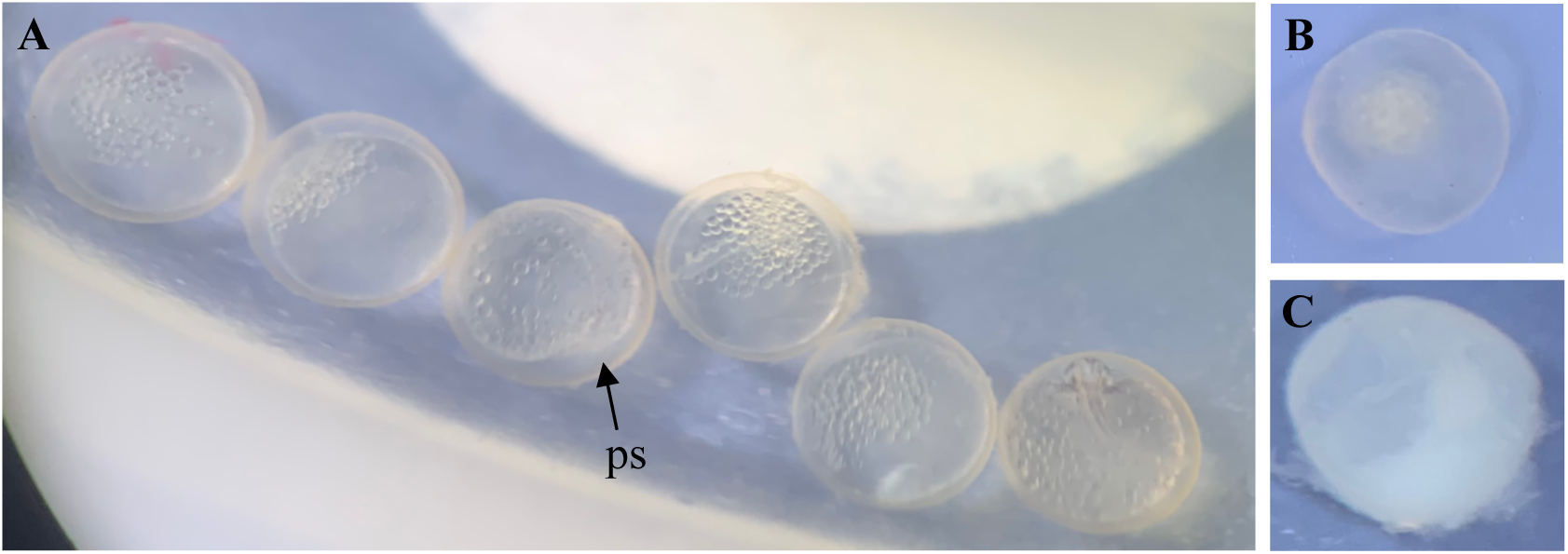
Scoring of eggs. Eggs were scored as A) fertilized, B) unfertilized, or C) inviable. Fertilized eggs were identified by the presence of the perivitelline space (ps). Unfertilized eggs lack the perivitelline space. Inviable eggs were identified by their opaque or discolored appearance.

### Fecundity and Self-compatibility Statistical Analysis

We examined the relationship between fecundity (number of eggs laid) and both the propensity for a given lineage to change sex, and the individual’s age. To determine whether fecundity varied as a function of the lineages’ propensities to change sex, we first ran a nominal logistic model with ‘laid eggs’ as a categorical dependent variable (yes = 1, no = 0) and lineage sex change propensity as the independent variable. We used a general linear model to examine the relationship between total number of eggs laid (continuous, dependent) and lineage sex change propensity (independent). Residuals of the total number of eggs were left skewed, so we transformed the number of eggs with square root (x+1). To determine whether fecundity (total number of eggs laid) varied as a function of age, we used a generalized regression model with a zero-inflated Poisson distribution and parent ID as a random effect to account for the fact that each parent was checked for eggs multiple times. An initial plot of the raw data suggested a possible non-linear relationship between age and fecundity so, we constructed linear, quadratic and cubic models and compared model fits using Akaike’s Information Criterion (AIC). The linear, quadratic and cubic age terms were scaled by mean and standard deviation (i.e., z-score) prior to running the models. The cubic model fit best (Table 3.1), so we report the results of this model.

We modeled self-compatibility in two ways: by the proportion of eggs that were fertilized and by hatch success of fertilized eggs. To determine whether the proportion of fertilized eggs varied as a function of the lineages’ propensity to change sex, we ran a general linear model. To determine whether the proportion of fertilized eggs varied as a function of the parents’ age when the eggs were collected, we ran a general linear model with parent ID as a random effect. To determine whether the proportion of fertilized eggs varied as a function of the total number of eggs laid by each parent, we ran a general linear model. We calculated hatch success as the proportion of fertilized eggs that resulted in a live hatchling. To determine whether hatch success varied as a function of the lineages’ propensity to change sex, we ran a general linear model. Models were run in JMP Pro version 15.0.0 (JMP®, Version 15 Pro 2019) and in RStudio using the glmmTMB package (RStudioTeam 2016; Brooks et al. 2017; R Core Team 2018). Data files will be deposited on GitHub upon publication acceptance.

## Results

The experiment included 213 individual fish from 100 different lineages. Thirty-five of those fish changed sex to male (16.4%) before the experiment ended at 365 - 371 dph. Of those 35, three individuals laid eggs first (9% of males, 1% of the total cohort). We collected 14,310 eggs, had 10,267 hatchlings (71.7% of eggs hatched), and 10,059 live hatchlings (98% of hatchlings were alive). The three individuals that changed sex to male laid a total of 69 eggs by 197 dph (the oldest age at which a fish was identified as male), which equated to, on average, 23 ± 13 (SEM) eggs per individual among the three, but 1.97 ± 1.4 eggs per capita for all individuals that changed sex to male. The remaining 178 hermaphrodites laid a total of 7,341 eggs before they collectively aged to 197 dph (41.2 ± 2.2 eggs per individual). We started checking for eggs from each fish at 67 dph and collected a single egg from one individual on the first egg check; the egg was fertilized. Overall, the proportion of eggs that were fertilized was 93.4%. The proportion of eggs that were unfertilized was 2.5%, and the proportion identified as “inviable” was 4.1%.

Our results indicate that fecundity initially increases with age until about 175 dph, then decreases until it plateaus at about 350 dph (Table 1 Figure 2A). The proportion of fertilized eggs also increased significantly with age, but not the total fecundity of the parent (Table 1, Figure 2B). Because we did not record the hatch success of each individual collection (i.e., clutch) of eggs, we were not able to analyze whether hatch success changed with age. The lineages’ propensity to change sex was significantly related to fecundity and self-compatibility. When individuals that changed sex to male were included in the model, the odds of laying any eggs decreased significantly as the propensity to change sex increased (Table 2). However, this relationship became non-significant when individuals that changed sex to male were excluded (Table 2). Conversely, the number of eggs laid decreased significantly as the propensity to change sex increased, whether or not males were included in the model (Table 2 Figure 3A). Both the proportion of fertilized eggs and the hatch success of those eggs also decreased significantly with increasing propensity to change sex (Table 2, Figure 3B, C).

**Figure 2.**
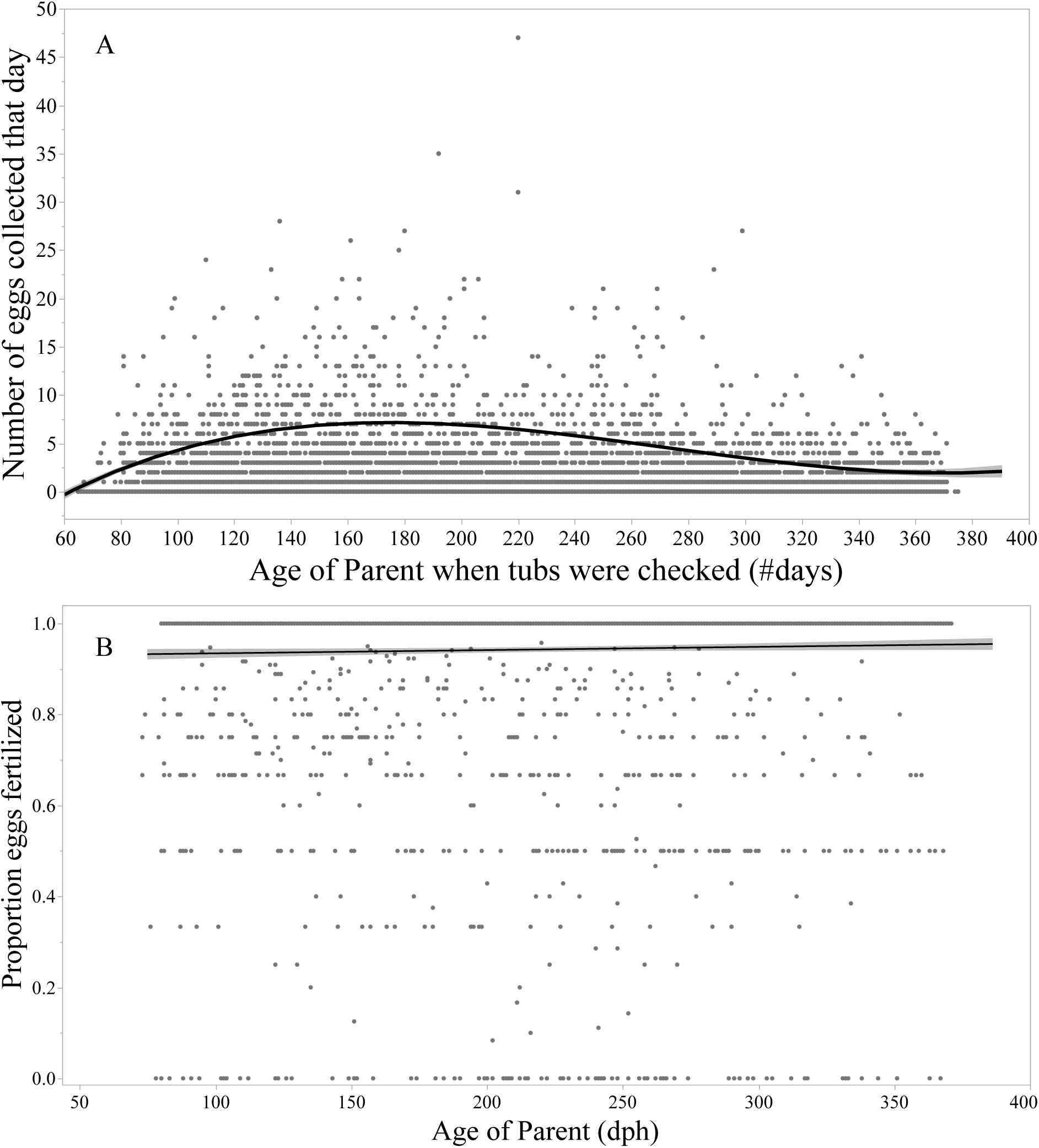
Age effects on the number of eggs collected and the proportion of those eggs that were fertilized. A) Age affected fecundity cubically; the number of eggs oviposited increased initially, then decreased between 170 – 180 dph, then plateaued around 350 dph. B) The proportion of eggs that were fertilized increased significantly with fish age. The additional straight line at the top of the graph is a string of points representing all of the egg collections where 100% of the eggs were fertilized.

**Figure 3.**
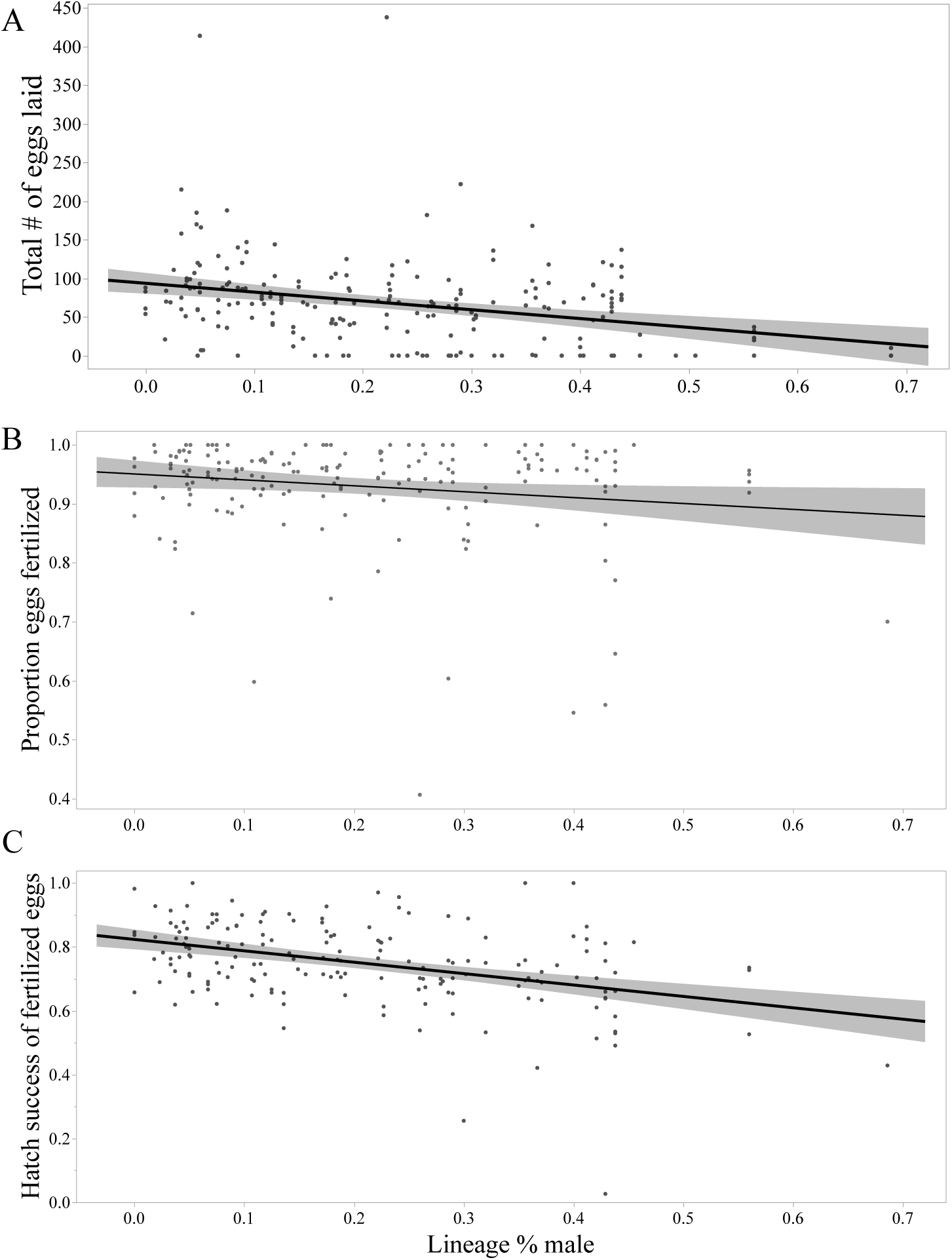
Lineage sex change propensity effects on A) fecundity, B) the proportion of eggs that were fertilized, and C) hatch success of fertilized eggs. Lineage sex change propensity is measured as the proportion of each lineage that change sex in isolation under common garden conditions in our colony (lineage % male).

**Table 1.** Summary of fecundity and self-compatibility models with age of parent fish as the dependent variable. AIC = Akaike’s Information Criterion; ZI = zero inflation; F = F-ratio; P = P-value; Z = Z-value; df = degrees of freedom.

| Fecundity |  |  |  |  |  |  |
| --- | --- | --- | --- | --- | --- | --- |
| Models | AICc | ΔAICc | ZI parameter<br>estimate<br>(p-value) | Estimate | Z | P |
| # eggs ~ _age + age<sup>2</sup><br>+ age<sup>3</sup> | 33140.0 | 0.0 | -0.98 (<0.0001) | age: 3.06<br>age<sup>2</sup>: -5.76<br>age<sup>3</sup>: 2.52 | 10.73<br>9.85<br>7.94 | <0.0001<br><0.0001<br><0.0001 |
| # eggs ~ _age + age<sup>2</sup> | 33198.5 | 58.5 | -0.92 (<0.0001) | age: 0.85<br>age<sup>2</sup>: -1.14 | 13.01<br>-17.67 | <0.0001<br><0.0001 |
| # eggs ~ _age | 33527.0 | 387.0 | -0.83 (<0.0001) | age: -0.29 | -29.58 | <0.0001 |
| Proportion of fertilized eggs |  |  |  |  |  |  |
|  |  |  | Estimate | F | df | P |
| Parent Age |  |  | 0.000069 | 4.08 | 1 | 0.04 |
| Number of Eggs Laid |  |  | 0.000093 | 0.63 | 1 | 0.43 |

**Table 2.** Summary of fecundity, self-compatibility, and hatch success models with lineage sex change propensity as the dependent variable. Fecundity was square root (x+1) transformed to ensure normality. The models for proportion of fertilized eggs and hatch success already excluded the males that did not lay any eggs, thus separate models were not needed. Significant effects are shown in bold. F = F-ratio, P = P-value, df = degrees of freedom.

| *Probability of Laying Eggs* | *Odds Ratio* | $\chi^2$ | *df* | *P* |
| --- | --- | --- | --- | --- |
| Lineage sex change propensity, **including** males | 0.003 | 23.85 | 1 | **<0.0001** |
| Lineage sex change propensity, **excluding** males | 0.006 | 2.62 | 1 | 0.11 |
| *Fecundity (# of eggs laid)* | *Estimate* | *F* | *df* | *P* |
| Lineage sex change propensity, **including** males | -4.17 | 36.29 | 1 | **<0.0001** |
| Lineage sex change propensity, **excluding** males | -4.25 | 9.09 | 1 | **0.003** |
| *Proportion of fertilized eggs* | *Estimate* | *F* | *df* | *P* |
| Lineage sex change propensity, **including** males | -0.10 | 4.86 | 1 | **0.029** |
| *Hatch Success* | *Estimate* | *F* | *df* | *P* |
| Lineage sex change propensity, **including** males | -0.36 | 33.85 | 1 | **<0.0001** |

## Discussion

Mixed mating strategies offer the fitness rewards of both outcrossing with conspecifics and fertilizing one’s own eggs. The cost-to-benefit ratio of each strategy is expected to vary across space and time as the external and internal environments also change (Maynard Smith 1971; Lively and Lloyd 1990; Lehtonen et al. 2012; Layman et al. 2017; Lynch et al. 2018). There is also variation among individuals in characters that affect mating strategy: fecundity and self-compatibility (Stephenson et al. 2000; Travers et al. 2004; Busch 2005). Variation in self-compatibility, combined with external factors such as pollinator and/or mate availability or parasite pressure, has the potential to drive evolutionary changes in mating strategy. For instance, if self-compatible individuals have greater reproductive fitness (produce more offspring and/or offspring have greater survival rates) and survive just as well as self-incompatible individuals, then the population may evolve to complete or near complete selfing. Comparatively, if there are plenty of mates/pollinators and/or inbreeding depression that increases the benefits to cost ratio of outcrossing, populations will evolve to complete or near complete outcrossing. We tested, in the only self-fertilizing hermaphroditic vertebrate, the hypothesis that self-compatibility and fecundity would vary with age and as a function of the lineages’ propensities to change sex.

The most important finding was that both fecundity and self-compatibility decreased as lineages became more likely to change sex, which is the exact opposite of what we predicted. We predicted that fecundity and/or self-compatibility would increase with the lineages’ propensity to change sex for one of two reasons: either individuals that eventually change sex would produce more eggs with greater self-compatibility, or the remaining hermaphrodites from high sex changing lineages would produce at least as many eggs with equal self-compatibility as those from lineages less likely to change sex. Given that neither of these predictions were supported, it remains unclear how sex change from a self-fertilizing hermaphrodite to an obligate outcrossing male is maintained in rivulus populations. Theory suggests that males incur a cost because, when compared to self-fertilizing hermaphrodites, they cannot produce offspring and also reduce the parent’s genetic contribution to offspring by one half (Maynard Smith 1971, 1978). Males must overcome this cost by producing many extremely fit offspring.

Prior experiments have provided evidence of outbreeding depression on fecundity (Gresham et al. 2020) in several ecologically relevant conditions, increasing the cost of males and outcrossing. However, Ellison et al. (2011) provide evidence of potential inbreeding depression under parasitic stress, therefore decreasing the cost of males. Male size also increases with heterozygosity, possibly allowing more heterozygous males to secure more resources and have greater access to unfertilized eggs (Molloy et al. 2011). The results from this experiment suggest that the cost of being male is highly variable among lineages. Only three individuals laid eggs before changing sex to male, indicating that they were not as fecund as self-fertilizing hermaphrodites. The odds of laying any eggs decreased as sex change propensity increased when all individuals were included in the model. However, when we excluded individuals that changed sex, there was not an effect of lineage sex change propensity, demonstrating that the hermaphrodites from high sex changing lineages are just as likely to lay *an* egg as those from low sex changing lineages. In contrast, hermaphrodites from high sex changing lineages laid significantly fewer eggs, even if the males were excluded from the model. Self-compatibility and hatch success also decreased as propensity to change sex increased. These results indicate that individuals from high sex changing lineages that remain hermaphrodite are not very fecund or self-compatible. This exemplifies that the cost of changing sex to male might vary among genetically distinct lineages. In particular, individuals from low sex changing, highly fecund and self-compatible lineages are likely to experience the highest cost of sex change.

We found that fecundity initially increases with age, then decreases, and finally plateaus as individuals age from about 60 dph to about 370 dph. The only other report of potential age-related changes in fecundity in rivulus comes from Harrington (1971). From 37 fish, he reported significant seasonality in “ovarian function” (i.e., fecundity) over three years when exposed to natural day light cycles, but not a significant change in fecundity with age. The seasonal decline reported by Harrington (1971), which occurred from September through December, aligns with the end of our experiment. The fish in our experiment hatched and then were removed from the experiment over a four-month period between 30 August 2018 and 22 December 2019. It is possible that the association between age and fecundity that we found, particularly the shallow decline in egg production from 180-300 dph represents the same seasonal decline described by Harrington (1971). He allowed his fish to experience natural daylight cycles. Our fish were kept on a constant photoperiod and a relatively constant temperature, and so should have been removed from any external signals of season. To truly understand how fecundity is affected by age and seasonality, we need additional years of records from multiple lineages.

Overall, 93.4% of the eggs we collected were fertilized. This is very close to the proportion of eggs fertilized reported by Harrington (1971). From six fish of one lineage and 1,056 eggs, he identified 94.3% as fertilized. We recorded 2.5% unfertilized and 4.1% inviable, compared to Harrington (1971), who reported 4% unfertilized and 1.7% inviable. It is likely that most of the inviable eggs we recorded were unfertilized when oviposited, but then became inviable and discolored by the time we collected them. Harrington (1971) collected eggs every day and was more likely to record unfertilized eggs before they became inviable. Our first fertilized egg was collected at 67 dph, indicating the egg was oviposited sometime between 60 dph and 67 dph. This is the earliest reported oviposition in rivulus.

Regardless of the cost of changing sex to male, there is still the issue of passing sex changing alleles to the next generation. How do lineages that are less fecund, less self-compatible, and that frequently change sex to male not disappear from populations? It is possible that many of these lineages do go extinct but, some Belizean populations maintain relatively high male proportions (> 20%) and non-trivial rates of outcrossing (Davis 1990; Turner et al. 1992; Tatarenkov et al. 2009, 2015), which suggests that males and outcrossing are under positive selection in some places. Males must be able to successfully find unfertilized eggs, but we are not sure how that happens. There have been multiple attempts to mate males and hermaphrodites in the lab. Mackiewicz et al. (2006a) paired multiple nearly-senescent hermaphrodites and males in individual tubs and confirmed two out of 32 offspring were outcrossed progeny. Nakamura et al. (2008) surgically removed eggs from two hermaphrodites for *in vitro* outcrossing, but only one out of thirteen hatchlings were outcrossed instead of self-fertilized. Our laboratory has tried pairing hermaphrodites and males in different ratios with no outcrossing success. In behavior trials, reported interactions between pairs of hermaphrodites and males have ranged from no courtship behaviors (Martin 2007) to significantly more courtship behaviors than same sex pairs (Luke and Bechler 2010) or courtship behaviors by the males with reciprocal attacks by the hermaphrodites (Turner et al. 1992). It is clear that we have not yet reproducibly identified conditions that favor outcrossing.

There often are opposing selection pressures for and against biparental sex (outcrossing). Mixed mating individuals can benefit from outcrossing when the costs of biparental sex are low, but self when the costs of biparental sex are high or when mates and/or pollinators are sparse. The relative benefit of selfing or outcrossing among individuals is also dependent on the individual level of self-compatibility, which can have a genetic basis and/or respond to the external environment in a plastic or flexible manner. Individuals and lineages that are not very self-compatible cannot exploit the benefits of selfing as much as individuals that are more self-compatible, even when the costs of outcrossing are high. In the mangrove rivulus fish, we have found significant variation in self-compatibility that correlates with age and with the lineages’ propensities to change sex. Individuals from lineages with a higher sex change propensity laid significantly fewer eggs and a significantly lower proportion of the eggs were self-fertilized (i.e., they were less self-compatible). Also, the proportion of fertilized eggs increased with age.

In wild populations, interactions between individuals’ level of self-compatibility and external conditions have the potential to drive evolutionary changes. The internal and external environments can alter the age structure of populations and/or the economics of outcrossing and selfing. In populations that have few or no lineages that frequently change sex (such as those recently founded by highly self-compatible individuals), lineages with higher propensities to change sex would likely struggle to become established, if offspring of selfed and outcrossed matings are equally fit. In these populations, individuals that change sex frequently will find an environment with very few unfertilized eggs, and very few opportunities to pass their sex changing alleles on to the next generation. However, once high sex change propensity lineages become established, possibly due to additional migration or outcrossed progeny having a fitness advantage over selfed progeny, these lineages can maintain sex change and males. Indeed, among rivulus populations across the geographic range we find a significant increase in the rate of outcrossing that correlates with an increase in the proportion of sex changing lineages (Mackiewicz et al. 2006b; Tatarenkov et al. 2015). We also find variation among lineages within in a population in the propensity to change sex, but the magnitude of variation is population specific (Gresham et al. 2020).

Males, and lineages that frequently produce males, may also have better success in populations with many young hermaphrodites. While we did not find evidence that younger hermaphrodites reproduce as “pure females” and only lay unfertilized eggs, younger hermaphrodites did lay more unfertilized eggs. If the fitness of progeny derived from selfing and outcrossing is relatively equal, population age structure can act as a selection pressure for or against lineages that frequently change sex. For instance, populations that lose a greater proportion of juveniles and young adults to predation will have proportionally fewer unfertilized eggs available for males than those populations in which younger individuals thrive; selecting against high sex changing lineages. These results open up fascinating opportunities for continued laboratory studies on lineage variation in traits such as size, fecundity, and sex change across different environments, and continued field studies to track demographic and population genetic characteristics such as selfing and outcrossing rates, the proportion of males, and population size and density.

## Notes

### Competing Interest Statement

The authors have declared no competing interest.

### Summary of Updates

added author and changed affiliation of corresponding author.

## References

Allard, R. W., G. R. Babbel, M. T. Clegg, and a L. Kahler. 1972. Evidence for coadaptation in *Avena barbata*. Proceedings of the National Academy of Sciences of the United States of America 69:3043–8.

Baker, H. 1955. Self-compatibility and establishment after “long-distance” dispersal. Evolution 9:347–349.

Bell, G. 1988. Recombination and the immortality of the germ line. Journal of Evolutionary Biology 1:67–82.

Brooks, M. E., K. Kristensen, K. J. van Benthem, A. Magnusson, C. W. Berg, A. Nielsen, H. J. Skaug, et al. 2017. glmmTMB balances speed and flexibility among packages for zero-inflated generalized linear mixed modeling. The R Journal 9:378–400.

Busch, J. W. 2005. The evolution of self-compatibility in geographically peripheral populations of *Leavenworthia alabamica* (Brassicaceae). American Journal of Botany 92:1503–1512.

Clutton-Brock, T. H., and G. A. Parker. 1995. Sexual coercion in animal societies. Animal Behaviour 49:1345–1365.

Cole, K. S., and D. L. G. Noakes. 1997. Gonadal development and sexual allocation in mangrove killifish, *Rivulus marmoratus* (Pisces: Atherinomorpha). Copeia 1997:596–600.

Darwin, C. 1876. The effects of cross and self fertilisation in the vegetable kingdom. D. Appleton.

Davis, W. P. 1990. Field observations of the ecology and habits of mangrove rivulus. Ichthyological Exploration of Freshwaters 1:123–134.

Dolgin, E. S., B. Charlesworth, S. E. Baird, and A. D. Cutter. 2007. Inbreeding and outbreeding depression in *Caenorhabditis* nematodes. Evolution 61:1339–1352.

Ellison, A., J. Cable, and S. Consuegra. 2011. Best of both worlds? association between outcrossing and parasite loads in a selfing fish. Evolution 65:3021–3026.

Escobar, J. S., J. R. Auld, A. C. Correa, J. M. Alonso, Y. K. Bony, M. A. Coutellec, J. M. Koene, et al. 2011. Patterns of mating-system evolution in hermaphroditic animals: Correlations among selfing rate, inbreeding depression, and the timing of reproduction. Evolution 65:1233–1253.

Fowler, K., and L. Partridge. 1989. A cost of mating in female fruitflies. Nature 338:760–761.

Furness, A. I., A. Tatarenkov, and J. C. Avise. 2015. A genetic test for whether pairs of hermaphrodites can cross-fertilize in a selfing killifish. Journal of Heredity 106:749–752.

Gresham, J. D., K. M. Marson, A. Tatarenkov, and R. L. Earley. 2020. Sex change as a survival strategy. Evolutionary Ecology 34:27–40.

Harrington Jr, R. W. 1971. How ecological and genetic factors interact to determine when self-fertlizing hermaphrodites of *Rivulus marmoratus* change into functional secondary males, with a reappraisal of the modes of intersexuality among fishes. Copeia 1971:389–432.

Hartfield, M., and P. D. Keightley. 2012. Current hypotheses for the evolution of sex and recombination. Integrative Zoology 7:192–209.

JMP®, Version 15 Pro. 2019. SAS Institute Inc., Cary, NC.

Layman, N. C., M. T. R. Fernando, C. R. Herlihy, and J. W. Busch. 2017. Costs of selfing prevent the spread of a self-compatibility mutation that causes reproductive assurance. Evolution 71:884–897.

Lehtonen, J., M. D. Jennions, and H. Kokko. 2012. The many costs of sex. Trends in Ecology & Evolution 27:172–178.

Lively, C. M. 2010. A review of Red Queen models for the persistence of obligate sexual reproduction. Journal of Heredity 101:S13–S20.

Lively, C. M., and D. G. Lloyd. 1990. The cost of biparental sex under individual selection. The American Naturalist 135:489–500.

Luke, K. N., and D. L. Bechler. 2010. The role of dyadic interactions in the mixed-mating strategies of the mangrove rivulus *Kryptolebias marmoratus*. Current Zoology 56:6–17.

Lynch, Z. R., M. J. Penley, and L. T. Morran. 2018. Turnover in local parasite populations temporarily favors host outcrossing over self-fertilization during experimental evolution. Ecology and Evolution.

Mackiewicz, M., A. Tatarenkov, A. Perry, J. R. Martin, J. F. Elder, D. L. Bechler, and J. C. Avise. 2006a. Microsatellite documentation of male-mediated outcrossing between inbred laboratory strains of the self-fertilizing mangrove killifish (*Kryptolebias marmoratus*). The Journal of Heredity 97:508–13.

Mackiewicz, M., A. Tatarenkov, D. S. Taylor, B. J. Turner, and J. C. Avise. 2006b. Extensive outcrossing and androdioecy in a vertebrate species that otherwise reproduces as a self-fertilizing hermaphrodite. Proceedings of the National Academy of Sciences of the United States of America 103:9924–8.

Martin, S. B. 2007. Association behaviour of the self-fertilizing *Kryptolebias marmoratus* (Poey): The influence of microhabitat use on the potential for a complex mating system. Journal of Fish Biology 71:1383–1392.

Maynard Smith, J. 1971. What use is sex? Journal of Theoretical Biology 30:319–335.

Maynard Smith, J. 1978. The evolution of sex. Cambridge Univ Press, Cambridge.

Molloy, P. P., E. A. Nyboer, and I. M. Côté. 2011. Male-Male Competition in a Mixed-Mating Fish. Ethology 117:586–596.

Morran, L. T., O. G. Schmidt, I. a Gelarden, R. C. Parrish, and C. M. Lively. 2011. Running with the Red Queen: host-parasite coevolution selects for biparental sex. Science (New York, N.Y.) 333:216–8.

Muller, H. J. 1932. Some genetic aspects of sex. American Society of Naturalists 703:118–138.

Nakamura, Y., K. Suga, Y. Sakakura, T. Sakamoto, and A. Hagiwara. 2008. Genetic and growth differences in the outcrossings between two clonal strains of the self-fertilizing mangrove killifish. Canadian Journal of Zoology 86:976–982.

Pannell, J. R., J. R. Auld, Y. Brandvain, M. Burd, J. W. Busch, P. Cheptou, J. K. Conner, et al. 2015. The scope of Baker’s law. New Phytologist 208:656–667.

R Core Team. 2018. R: A language and environment for statistical computing. R Foundation for Statistical Computing, Vienna, Austria.

RStudioTeam. 2016. RStudio: Integrated Development for R. RStudio, Inc., Boston.

Scarsella, G. E., J. D. Gresham, and R. L. Earley. 2018. Relationships between external sexually dimorphic characteristics and internal gonadal morphology in a sex-changing fish. Journal of Zoology 305:133–140.

Stebbins, G. L. 1957. Self fertilization and population variability in the higher plants. The American Naturalist 91:337.

Stephenson, A. G., S. V Good, and D. W. Vogler. 2000. Interrelationships among inbreeding depression, plasticity in the self-incompatibility system, and the breeding system of *Campanula rapunculoides* L. (Campanulaceae). Annals of Botany 85:211–219.

Tatarenkov, A., R. L. Earley, B. M. Perlman, D. S. Taylor, B. J. Turner, and J. C. Avise. 2015. Genetic subdivision and variation in selfing rates among Central American populations of the mangrove rivulus, *Kryptolebias marmoratus*. Journal of Heredity 106:276–284.

Tatarenkov, A., S. M. Q. Lima, D. S. Taylor, and J. C. Avise. 2009. Long-term retention of self-fertilization in a fish clade. Proceedings of the National Academy of Sciences of the United States of America 106:14456–14459.

Tatarenkov, A., B. C. Ring, J. F. Elder, D. L. Bechler, and J. C. Avise. 2010. Genetic composition of laboratory stocks of the self-fertilizing fish *Kryptolebias marmoratus*: a valuable resource for experimental research. PloS one 5:e12863.

Travers, S. E., J. Mena-Ali, and A. G. Stephenson. 2004. Plasticity in the self-incompatibility system of *Solanum carolinense*. Plant Species Biology 19:127–135.

Tsitrone, A., P. Jarne, and P. David. 2003. Delayed Selfing and Resource Reallocations in Relation to Mate Availability in the Freshwater Snail *Physa acuta*. American Naturalist 162:474–488.

Turner, B. J., W. P. Davis, and D. S. Taylor. 1992. Abundant males in populations of a selfing hermaphrodite fish, *Rivulus marmoratus*, from some Belize cays. Journal of Fish Biology 40:307–310.

Winn, A. A., E. Elle, S. Kalisz, P. Cheptou, C. G. Eckert, C. Goodwillie, M. O. Johnston, et al. 2011. Analysis of inbreeding depression in mixed-mating plants provides evidence for selective interference and stable mixed mating. Evolution 65:3339–3359.

Wright, S. I., S. Kalisz, and T. Slotte. 2013. Evolutionary consequences of self-fertilization in plants. Proceedings. Biological sciences / The Royal Society 280:20130133.

